# Comparative analysis of zooplankton diversity in freshwaters: What can we gain from metagenomic analysis?

**DOI:** 10.1101/2021.10.27.465999

**Authors:** Marie-Ève Monchamp, David A. Walsh, Rebecca E. Garner, Susanne A. Kraemer, Beatrix E. Beisner, Melania E. Cristescu, Irene Gregory-Eaves

**Affiliations:** Department of Biology, McGill University, Montreal, Quebec, Canada; Groupe de recherche interuniversitaire en limnologie (GRIL); Department of Biology, Concordia University, Montreal, Quebec, Canada; Department of Biological Sciences, University of Quebec at Montreal, Montreal, Quebec, Canada

**Keywords:** Shotgun sequencing, Environmental DNA (eDNA), Freshwater ecology, Biodiversity survey, Ecological assessment

## Abstract

Molecular genetic approaches applied to environmental DNA have great potential for biodiversity research and ecosystem monitoring. A metagenome contains genetic information from all organisms captured in an environmental sample. It has been primarily used to study bacteria and archaea, but promising reports focusing on metazoan diversity are emerging. However, methodological uncertainties remain, and studies are required to validate the power and the limitations of such an approach when applied to macro-eukaryotes. Here, we analyzed water sample metagenomes to estimate zooplankton diversity in 22 freshwater lakes across Eastern Canada. We tested the coherence of data based on morphologically identified zooplankton taxa and molecular genetic data derived from shotgun sequencing of environmental DNA collected at the same time. RV coefficients showed a significant correlation between the relative abundance of zooplankton families derived from small subunit rRNA genes extracted from the metagenomes and morphologically identified zooplankton. However, differences in congruence with morphological counts were detected when varied bioinformatic approaches were applied to presence-absence data. This study presents one of the first diversity assessments of a group of aquatic metazoans using metagenomes and validates the coherence of the community composition derived from genetic and classical species surveys. Overall, our results suggest that metagenomics has the potential to be further developed to describe metazoan biodiversity in aquatic ecosystems, and to advance this area we provide key recommendations for workflow improvement.

## 1. Introduction

In the context of intensifying global change, there is a growing need for broad scale monitoring strategies and ecosystem assessment (Cardinale et al., 2012; Cordier et al., 2020). Approaches based on environmental DNA (eDNA), broadly defined as the total pool of DNA that can be isolated from the environment (Taberlet et al., 2012; Pawlowski, 2020; Rodriguez-Ezpeleta et al., 2021), represent high-throughput, cost-effective, non-invasive tools that are being increasingly used in biodiversity monitoring programs (Bohmann et al., 2014; Deiner et al., 2017). One of the most common methods to interpret the eDNA signal from a complex community is marker gene metabarcoding, which allows for multiple taxa to be investigated in a single sequencing experiment (Hajibabaei et al., 2011; Taberlet et al., 2012). This approach has led to numerous successful biodiversity assessments of terrestrial and aquatic biota, including metazoans (e.g. Hänfling et al., 2016; Sigsgaard et al., 2016; Deiner et al., 2017; Taberlet et al., 2018), and has the potential to help us gain a more holistic view of an ecosystem with hundreds of organisms identified simultaneously from one environmental sample. Metabarcoding is a highly sensitive approach that can detect rare or cryptic species (Thomsen et al., 2012; Port et al., 2016), and is seen as a promising approach in ecological assessment studies of aquatic ecosystems (Aylagas et al., 2016; Cordier et al., 2017; Yang and Zhang, 2020). Nevertheless, metabarcoding as well as other PCR-based techniques, such as quantitative PCR, introduce biases. For example, universal primers used to barcode multiple groups of taxa simultaneously do not necessarily bind equally to different templates, leading to amplification bias or the complete loss of certain groups (Tedersoo et al., 2015; Alberdi et al., 2018; Kelly et al., 2019).

Metagenomics, broadly defined as the application of high-throughput shotgun sequencing technologies to capture the entire pool of species present in an eDNA sample without targeting a specific gene marker (Tringe and Rubin, 2005), is an emerging approach but to date has been primarily applied to study microbial communities (Grossart et al., 2020). Metagenomics has only recently gained traction in the study of larger organisms such as metazoans and is now seen as a complement (Singer et al., 2020) and potential alternative to metabarcoding. There are several reasons that might explain the low number of studies using metagenomics to investigate eukaryotes (Barnes and Turner, 2016). First, the efficiency of metagenomics to capture the macro-eukaryote signal is not fully understood. Generally, it is believed that macro-eukaryotes are present in much lower densities in the environment compared to microbes (Azam and Malfatti, 2007), which might limit the recovery of the macro-eukaryote DNA signal. Second, issues related to the large size and low coding density of eukaryote nuclear genomes may contribute to poor recovery of eukaryotes in environmental metagenomes. For example, genomes of eukaryotes contain many repetitive elements that are difficult to assemble into scaffolds, as well as long non-coding sequences which are generally less taxonomically informative (Bik et al., 2012). Abundance estimations of eukaryotes based on shotgun sequencing are further complicated by the high interspecific variability in the number of rRNA gene copies per nucleus (Bik et al., 2012). Finally, both micro- and macro-organisms will often not find a match in reference databases unless they closely relate to an organism that has had its whole genome sequenced. This is a well-known challenge in any eDNA assessments, but curated DNA reference databases are growing rapidly, and thus it is expected that such limitations will continue to decrease in the near future (e.g., Lewin et al. 2018).

Despite the challenges, a handful of studies have shown promising results in applying metagenomics for broad biodiversity assessments of metazoans in water (e.g. Cowart et al. (2018); Singer et al.(2020); Machida et al. (2021); Manu et al. (2021)) and sediment samples (e.g. Pedersen et al. (2016); Gelabert et al. (2021)). This type of work, however, requires adapting bioinformatics pipelines to accommodate the diluted metazoan signal in eDNA, especially when targeting rare organisms. For instance, in microbial metagenomic studies, reads are typically assembled before being mapped to genomes for annotation. However, this approach is not always feasible when working with extra-organismal eDNA, likely due to the degraded nature and limited amount of starting genetic material (Barnes and Turner, 2016). An alternative to assembly is to annotate directly via mapping of metagenomic reads to reference databases of nucleotide or proteins sequences. Although computationally intensive, this approach has been reported effective when the output is processed using a Last Common Ancestor (LCA) algorithm or in combination with compositional interpolated Markov models (Quince et al., 2017). Given the relatively nascent nature of this field, we chose to evaluate the differences in diversity detected between targeting a taxonomically informative gene marker, the 18S rRNA gene in eukaryotes (i.e. SSU rRNA gene approach), vs. a broader analysis of the tens of millions of metagenomic reads (i.e. whole metagenome approach).

Here, we provide new insight on the effectiveness and reliability of metagenomics applied to extra-organismal eDNA in water samples for describing freshwater zooplankton. Our main questions are: i) can we effectively detect zooplankton diversity in lake water metagenomes, ii) how does the metagenomic gene prediction approach based on a single taxonomic marker (SSU rRNA gene) compare to mapping the entire eukaryotic fraction of metagenome reads, and iii) do diversity metrics derived from metagenomes show similar responses to key environmental gradients as those detected with morphological taxonomic surveys? We assessed zooplankton (Cladocera, Copepoda, and Rotifera) diversity based on surface water metagenomes from 22 lakes in Eastern Canada and compared these results with zooplankton data from morphologically identified samples collected in net hauls from the same sites. Our study is a timely response to the growing interest in adapting metagenomics techniques for advancing a holistic perspective of aquatic food webs across all domains of life from a single environmental snapshot.

## 2. Methods

### 2.1. Sites description

The 22 lakes were sampled as part of the Natural Sciences and Engineering Research Council of Canada (NSERC) Canadian Lake Pulse Network campaign in summer 2017 (Huot et al., 2019). (**Figure 1**). Lakes span a range of morphological characteristics and trophic status, as summarized in **Table S1**. Sampling occurred at a station situated at the maximum depth of each lake. The complete field protocol details are provided by LakePulse (NSERC Canadian Lake Pulse Network, 2021).

**Figure 1.**
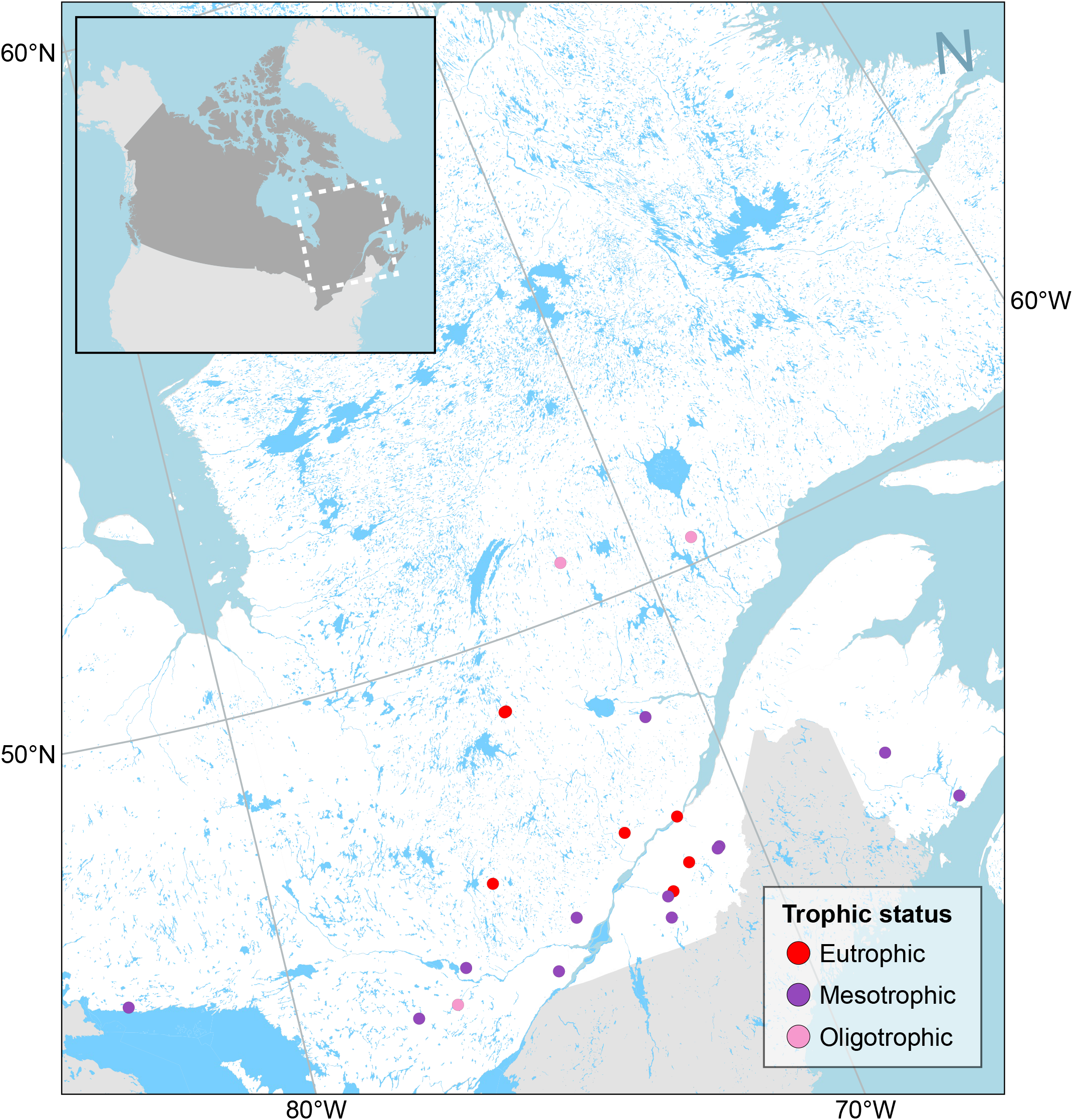
Map of Canada showing the location of the 22 lakes in Eastern Canada and their trophic status based on total phosphorus concentration: oligotrophic (less than 10 μg/L), mesotrophic (10 – 30 μg/L) and eutrophic (greater than 30 μg/L).

### 2.2. Sampling, DNA isolation, and taxonomic identification

#### 2.2.1. Zooplankton morphological identification

Crustacean zooplankton were sampled over the depth of the water column from 1 m above the sediment up to the water surface using a Wisconsin net with 100 μm mesh (10 cm net radius and 100 cm length). For relatively shallow lakes (<6 m-deep), additional vertical hauls were taken in the same manner to increase sample volume. Crustacean zooplankton were anesthetized with CO_2_ (Alka-Seltzer) and preserved in 70% ethanol (approx. final concentration) at room temperature. Species-level identification of crustacean zooplankton was done with a dissecting microscope under 100x to 400x magnification by BSA Environmental Services (Ohio, U.S.A.). Species biomass was estimated following the method from McCauley (1984). A detailed identification protocol is available in Paquette et al. (2021).

Rotifer counts were done on Lugol-preserved tow haul samples collected in the same manner as the crustacean zooplankton samples (above), except that instead of sampling from 1 m above the sediment to the lake surface, the rotifer samples were collected from the euphotic zone only. In several instances, the euphotic zone is identical to max depth minus 1 m. The coherence between the original cladoceran zooplankton counts and the rotifer counts performed on a different set of samples was verified by counting *Bosminidae* individuals in both sample types, to confirm that the preserved samples for rotifer counting were representative of the original zooplankton samples (**Supplementary File 1**).

#### 2.2.3. Environmental DNA sampling for metagenomic analyses

Water for eDNA was collected at the same station as the net hauls with an acid-washed integrated depth sampler over the euphotic zone down to 2 m below the surface. Our eDNA sampling strategy aimed at targeting mainly extra-organismal DNA, i.e. DNA that is not contained within whole organisms (Rodriguez-Ezpeleta et al., 2021), sometimes also referred to as ‘extracellular DNA’ (Taberlet et al., 2012; Bohmann et al., 2014). Thus, for samples dedicated to eDNA analysis, water was first passed through a 100 μm nylon mesh to remove large particles, and then up to 500 mL of water was vacuum-filtered on a Durapore 0.22 μm membrane (Sigma-Aldrich, St. Louis, USA) through a glass funnel apparatus at a maximum pressure of 8 inHg until the filter clogged. Filtrations were done on site in a tent, and filters were preserved immediately thereafter in cryovials at −80°C until analysis. Caution was taken to limit foreign DNA contamination in the field. All materials and equipment were acid-washed between lakes, and gloves were worn during sampling and filtering. In the laboratory, DNA was extracted from filters using the DNeasy PowerWater kit (QIAGEN, Hilden, Germany) following the manufacturer’s protocol with the addition of two steps as detailed by Garner et al. (2020). DNA was quantified using a Qubit 2.0 fluorometer and the dsDNA BR Assay kit (Invitrogen, Carlsbad, CA, USA). An aliquot of each DNA extract was sent to Genome Quebec facilities (Montreal, Canada) for shotgun library preparation and sequencing on an Illumina NovaSeq 6000 S4 PE150 with flow cell type S2.

### 2.3. Metagenomic analysis pipelines

Raw demultiplexed shotgun sequence files were quality checked using FastQC v.0.11.15. Adapter trimming and quality filtering were done with Trimmomatic v.0.38 (Bolger et al., 2014) with a minimum average quality of 25 and a minimum length of 36 nucleotides. We applied two slightly different approaches to identify eukaryote sequences in the metagenomes (**Figure 2**).

**Figure 2.**
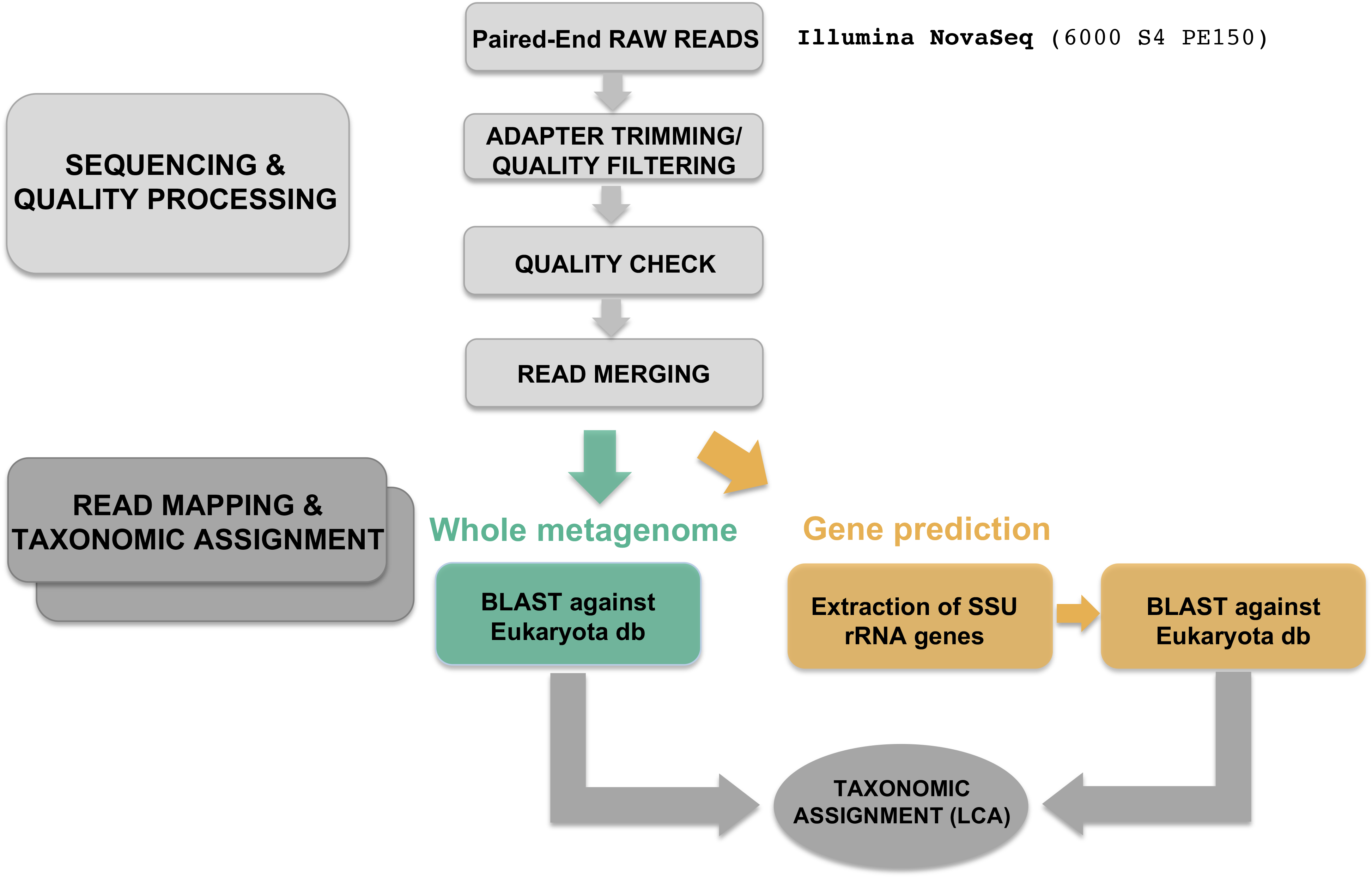
Detailed bioinformatics workflow used on shotgun sequencing data.

In the whole metagenome approach, all cleaned shotgun paired-end sequences were merged using PEAR (Zhang et al., 2014) before they were aligned against a local database consisting of all Eukarya entries in the NCBI non-redundant nucleotide database with the following parameters: min e-value 0.001, min percentage identity = 70, and retaining max 30 hits per read. BLASTn output files were then imported in MEGAN6 v.6.20.17 (Huson et al., 2016) for taxonomic assignment based on the lowest common ancestor (LCA) algorithm with a minimum score of 80, a minimum similarity of 80%, a minimum support of 2 reads and a minimum complexity filter set at 0.1. A detailed bioinformatic workflow is available as supplementary material (**Supplementary File 2**).

In the SSU rRNA gene prediction approach (corresponding to 18S rRNA genes in Metazoa), we applied the results of the ‘raw reads analysis pipeline’ of the European Bioinformatics Institute (EBI) MGnify (Mitchell et al., 2020). The detailed pipeline is described on the EBI website (https://emg-docs.readthedocs.io/en/latest/analysis.html#raw-reads-analysis-pipeline). Briefly, paired end reads were merged prior to adapter trimming and quality filtering. Additional non-coding RNAs (ncRNAs) were identified with Infernal (Nawrocki and Eddy, 2013) (HHM-only mode) using a library of ribosomal RNA hidden Markov models from Rfam (Kalvari et al., 2018) to identify large and small (LSU and SSU) rRNA genes. Following this, the reads identified as SSU rRNA genes were aligned with BLASTn and annotated following the whole metagenome approach described above (**Figure 2).**

### 2.4. Diversity analysis

Diversity analyses based on zooplankton assemblages surveyed using both microscopy and metagenomics were conducted in R v.4.1.0. (R Core Team, 2020). All diversity indices were calculated on assemblages binned to the family rank to deal with uneven taxonomic assignment resolution for different zooplankton groups across analytical platforms. The most common diversity metrics (taxonomic richness, Shannon index, Pielou’s evenness) were estimated on zooplankton abundance data (i.e. the number of individuals per liter or the number of sequencing reads) using the *diversity* function of the package vegan (Oksanen et al., 2013) and were used in least-square regressions against key environmental gradients identified from an earlier analysis of eastern Canadian LakePulse sites (Griffiths et al., 2021): epilimnetic total phosphorus concentration, specific conductivity, lake depth and an index of watershed disturbance calculated as the human impact index (HI) (Huot et al., 2019). All environmental variables were logarithm transformed, except for HI values (percentages) that were arcsine transformed.

Principal Component Analyses (PCA) were performed separately for each dataset using the function *prcomp* on both logarithm and Hellinger-transformed abundance (i.e. the number of individuals per liter or the number of reads sequenced) and biomass data where data were available (i.e. only for crustacean zooplankton observations) (Legendre and Gallagher, 2001). The three main principal components were extracted and used to derive an RV coefficient, analogous to Pearson’s correlation coefficient for two given multivariate data matrices (Legendre and Birks, 2012). All possible pairwise comparisons between datasets were explored – densities or biomass vs. either SSU rRNA genes or whole metagenome, and SSU rRNA genes vs. whole metagenome. Coefficient significance was verified with the function *coeffRV* in FactoMineR (Lê et al., 2008). We also considered the congruence between community identifications done for each sample using morphological data and shotgun analyses by calculating pairwise Jaccard and Bray-Curtis dissimilarities (the former based on incidence data and the latter based on relative abundance data (number of individuals per liter) using the function *vegdist* in vegan (Oksanen et al., 2013). For this analysis, no biomass data was used.

## 3. Results

### 3.1. Zooplankton taxonomy diversity across analytical platforms

Based on the microscopic analyses, we detected an average zooplankton family-level richness of 11.1 across the 18 lakes with complete zooplankton counts (**Table 1**; rotifer data were missing for three lakes). The most dominant families in terms of counts were the *Bosminidae*, *Cyclopidae* and *Daphniidae*, whereas the dominant families in terms of biomass (crustacean zooplankton only) were *Daphniidae*, *Cyclopidae*, and *Diaptomidae*. Across the 22 sites, the crustacean community was relatively even based on abundance data, with a Pielou’s evenness index of 0.67 (0 = no evenness, 1 = complete evenness). Considering just the crustacean zooplankton families for which there is a larger data set of hundreds of lakes across our four focal ecozones (Paquette et al., 2021), we found a comparable amount of richness and evenness within the range of the key environmental gradients captured by our 22 sites (**Table 1**).

**Table 1.** Summary of the main diversity indices (mean (min – max)) estimated for the three survey approaches at the Family taxonomic level. Microscopic estimates for Shannon and Pielou were based on abundance data (the number of individuals per liter, or the number of sequenced reads).

| **Approach** | **Taxonomic richness** | **Shannon index** | **Pielou's evenness** |
| --- | --- | --- | --- |
| Microscopy | 11.1 (7 – 15) | 1.6 (0.7 – 1.9) | 0.67 |
| SSU rRNA gene subset | 5.6 (1 – 10) | 1.2 (0 – 1.9) | 0.74 |
| Whole metagenome | 14.2 (10 – 21) | 1.3 (0 – 2.0) | 0.47 |
| Microscopy (198 LakePulse sites) | 6.49 (2 – 10) | 1.14 (0.1 – 1.8) | 0.61 |

High-throughput sequencing yielded on average ~28 million raw reads per metagenome and the number of reads per sample after quality filtering and merging of the pairs varied between 7 and 29 million **(Supplementary Figure S2**). Overall, the proportion of the merged reads assigned to Eukaryotes ranged between 0.5 and 1.2% of the total paired reads, with up to 46% of the eukaryotic reads confidently assigned to Metazoans (**Supplementary Figure S2**). With the whole metagenome BLAST approach, we detected a slightly greater average family richness of 15.95 (**Table 1**). Relative to the microscopy dataset we found that the assemblages in our 22 lakes were less even (mean Pielou’s evenness = 0.47; **Table 1**). The dominant taxa in terms of reads were *Daphniidae*, *Diaptomidae* and *Brachionidae* (Rotifera). Using the targeted SSU rRNA gene prediction approach, we detected the lowest average family richness relative to the previous two analytical approaches, with a mean of 5.6 (min = 1, max = 10). The dominant taxa detected were *Diaptomidae*, *Synchaetidae* (Rotifera) and *Cyclopidae*. Comparing across the different platforms, we found that the SSU rRNA gene prediction approach yielded the lowest family diversity values but evenness estimates that were closer to those generated through the microscopic counts for densities (**Table 1, Supplementary figure S3).**

### 3.2. Congruence of morphological and sequencing zooplankton families

We found a nested group of family diversity as we moved from SSU rRNA genes, to microscopy to whole metagenome datasets (**Figure 3a**). Nineteen out of 23 families that were detected at most sites using the whole metagenome approach were also found in the microscopy dataset of the 22 lakes. Families that were absent in the microscopy but present in the metagenomes are taxa that are often characterized as benthic or littoral associated (i.e. *Harpacticidae* (copepoda), *Chironomidae* (Diptera larvae), *Adinetidae* (Rotifera), and *Philodinidae* (Rotifera)).

**Figure 3.**
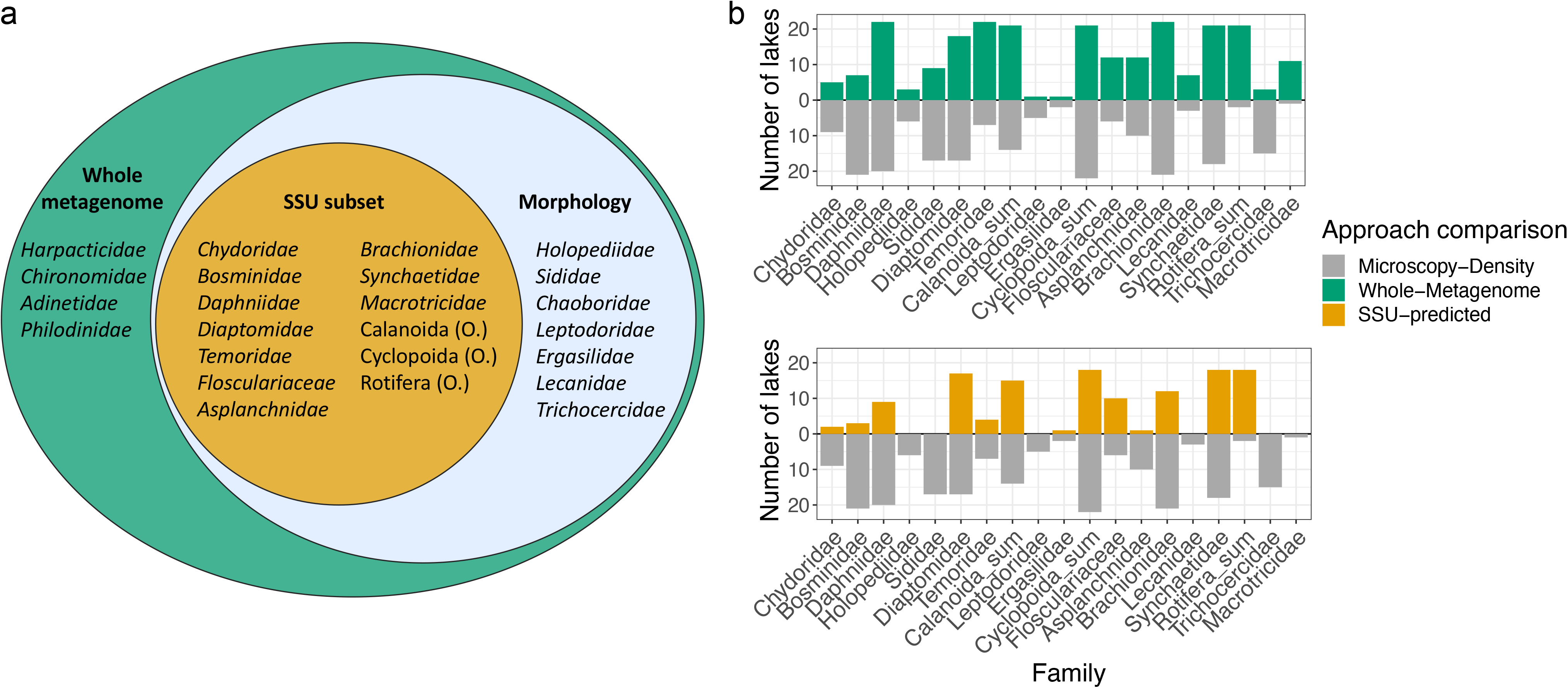
a) Diagram showing the number of zooplankton families and their overlap in detection via the three approaches: morphology-based microscopy (blue), whole metagenome sequencing (green), and gene prediction approach based on the small subunit subset of reads (orange). b) Divergent plots showing the number of lakes in which each zooplankton family were detected via the different approaches (grey: microscopic identification; green: whole-metagenome approach; orange: small-subunit (SSU) rRNA genes approach).

Zooplankton family occurrences across lakes were compared between the three analytical platforms (microscopy and two metagenomics approaches) to determine the level of congruence between survey methods (**Figure 3b & 4**). The families *Ergasilidae* (copepoda), *Leptodoridae* (cladocera), and *Holopediidae* (cladocera) were consistently absent at most sites (found only in a single or a few sites), whereas the Calanoida group (copepods - order level; found at 11 sites), *Synchaetidae* (rotifer; found at 11 sites), and the Cyclopoida group (copepods - order level; found at 13 of the 19 sites) were the three taxa that were most consistently widely detected across all analytical platforms (**Figure 4**). It is worth noting, however, that since the Calanoida and Cyclopoida groups were binned at order level, they are likely to comprise more than one Family each. The reason for this grouping was two-fold: firstly, the majority of the genetic reference sequences for these clades were lacking finer taxonomic resolution, and secondly, these groups include nauplii or juvenile stages which could not be assigned to one or the other order in the microscopy data based on morphological observations only.

**Figure 4.**
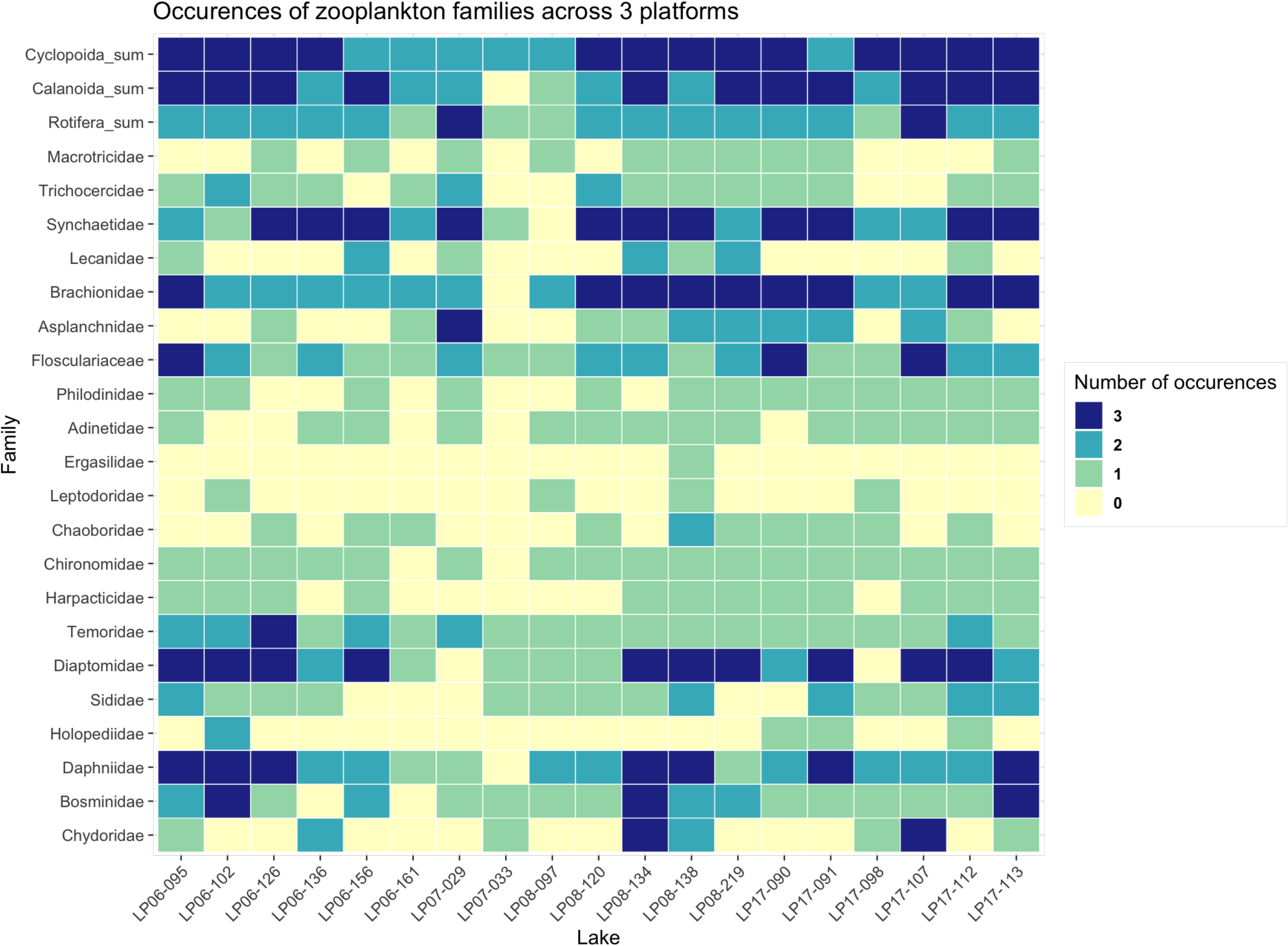
Heatmap with the number of each zooplankton Family-level (or Order-level in the case of Cyclopoida and Calanoida copepods and unidentified families of Rotifera) detection across analytical platforms (microscopy, whole metagenome, and small subunit (SSU) rRNA genes). A value of zero signifies that a Family/Order was absent from all datasets.

When comparing pairwise taxon occurrences across the three datasets for all zooplankton families and with copepods grouped at order level (Calanoida and Cyclopoida), we found consistent detections in 45% of cases (either 3 out of 3 or 0 out of 3 detections). When comparing microscopy with either genetic approach, the overall number of dual positive detections was higher between microscopy and whole metagenome datasets, with a total of 34.3% positive matches across 17 lakes (two lakes missing whole metagenome data were excluded) compared to only 17.3% positive matches for the comparison with SSU rRNA gene data in 19 lakes (**Figure 4**).

To consider the congruence of the entire assemblage between analytical platforms we calculated dissimilarity indices and RV coefficients. From a taxonomic presence-absence perspective (Jaccard distances), whole metagenomes more strongly reflected the community composition based on zooplankton counts than the SSU rRNA gene data (*p* = 0.0008). In contrast, when zooplankton family relative abundances (Bray-Curtis dissimilarities) were considered, we found that the SSU rRNA gene data performed similarly to the whole genome BLAST (*p* = 0.8051; **Figure 5**).

**Figure 5.**
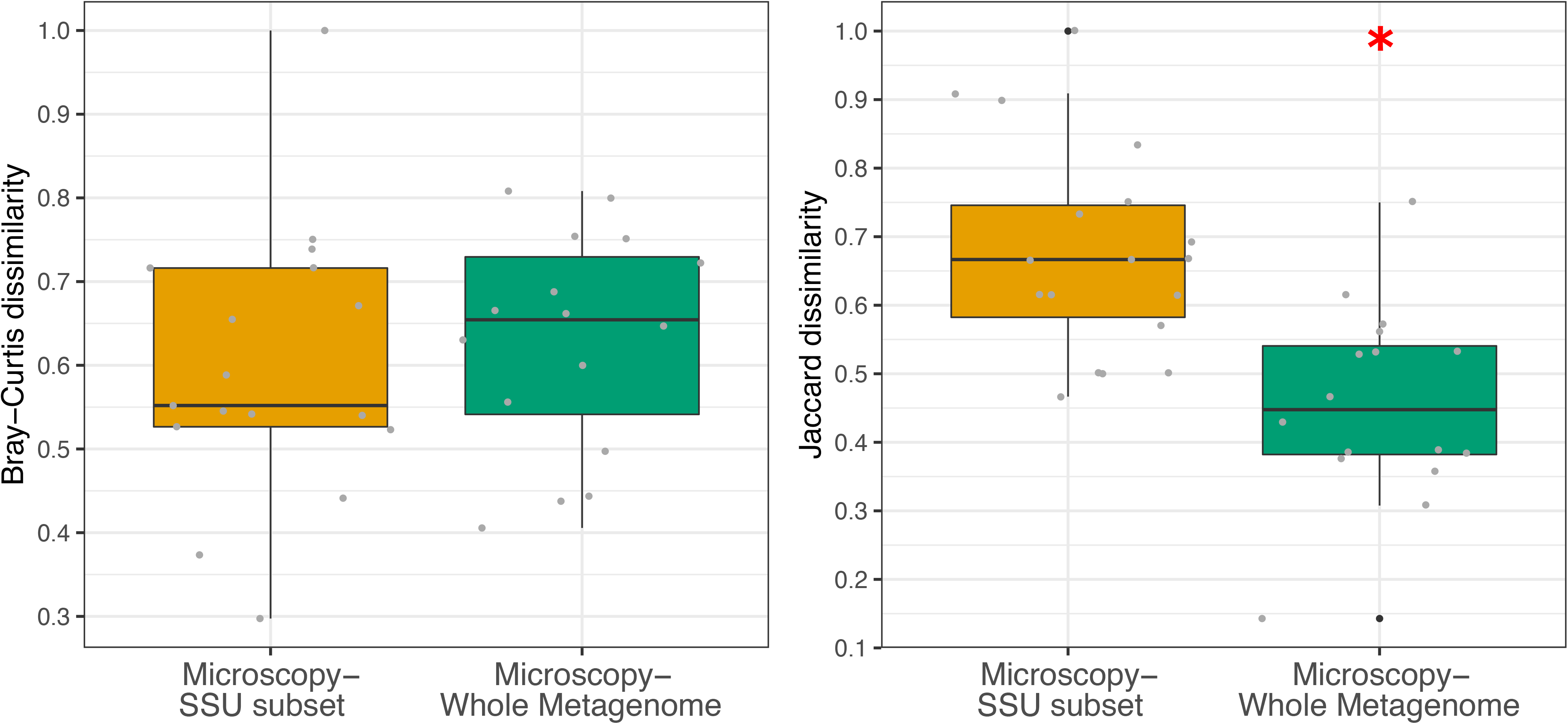
Boxplots showing Bray-Curtis (left) and Jaccard (right) dissimilarities between the microscopy-based taxonomic composition and the sequence read composition using the whole-metagenome approach (green) and the SSU rRNA gene subset approach (orange). The significant ANOVA test is indicated with an asterisk.

To further explore the congruence between assemblages, we calculated the RV coefficient based on the three main PCA axes and found moderate and significant congruence between SSU rRNA gene and morphology data (**Table 2**). Results were stronger when relative abundance data were log-transformed to account for potential over-representation of model organisms in the reference databases, which can yield inflated assigned read estimates compared to other taxonomic groups that are poorly populated in reference databases. When data were Hellinger-transformed, the RV coefficient between whole metagenome and SSU rRNA gene datasets was also of moderate strength (**Table 2**).

**Table 2.** Results of the congruence test between datasets (RV coefficients with significance values) using two types of data transformation (Hellinger or log_10_+1) on genetic and morphological data (abundance of all zooplankton and biomass of crustacean zooplankton only).

| Matrices compared | Taxa included | Transformation | RV coefficient | p-value |
| --- | --- | --- | --- | --- |
| Density-SSU | All zooplankton | Hellinger | **0.39** | **0.004** |
| Density-WM | All zooplankton | Hellinger | 0.006 | 0.93 |
| SSU-WM | All zooplankton | Hellinger | **0.38** | **0.003** |
| Density-SSU | All zooplankton | $\log_{10}+1$ | **0.48** | **0.0006** |
| Density-WM | All zooplankton | $\log_{10}+1$ | 0.12 | 0.39 |
| SSU-WM | All zooplankton | $\log_{10}+1$ | 0.17 | 0.19 |
| Biomass-SSU | Crustaceans only | Hellinger | 0.21 | 0.12 |
| Biomass-WM | Crustaceans only | Hellinger | 0.03 | 0.96 |
| Biomass-Density | Crustaceans only | Hellinger | **0.64** | **3.33E-06** |
| SSU-WM | Crustaceans only | Hellinger | **0.25** | **0.045** |
| Biomass-SSU | Crustaceans only | $\log_{10}+1$ | NS | NS |
| Biomass-WM | Crustaceans only | $\log_{10}+1$ | NS | NS |
| Biomass-Density | Crustaceans only | $\log_{10}+1$ | **0.51** | **0.0002** |
| SSU-WM | Crustaceans only | $\log_{10}+1$ | 0.12 | 0.28 |
| Density-SSU | Rotifers only | Hellinger | 0.09 | 0.68 |
| Density-WM | Rotifers only | Hellinger | 0.13 | 0.53 |
| SSU-WM | Rotifers only | Hellinger | 0.09 | 0.70 |
| Density-SSU | Rotifers only | $\log_{10}+1$ | 0.08 | 0.96 |
| Density-WM | Rotifers only | $\log_{10}+1$ | 0.10 | 0.66 |
| SSU-WM | Rotifers only | $\log_{10}+1$ | 0.02 | 0.96 |

### 3.3. Zooplankton diversity patterns over environmental gradients

We investigated the relationship between Family-level zooplankton diversity metrics (richness and Shannon diversity) and major environmental gradients identified in the eastern ecozones: the human impact index, total phosphorus (TP), specific conductivity and lake depth). Our analyses showed that several consistent relationships were apparent across the analytical platforms. For richness, significant negative relationships were observed between TP and both metagenomic datasets (SSU: adj. R^2^ = 0.24, *p* = 0.02; WM: adj. R^2^ = 0.29, *p* = 0.01) (**Figure 6 & Supplementary Table S2**). A negative but non-significant relationship (*p* = 0.24) was observed between TP and richness derived from morphological identifications. We also detected negative relationships between richness and human impact index, but the fit once again was only significant for the metagenomic datasets (SSU: adj. R^2^ = 0.23, *p* = 0.022; WM: adj. R^2^=0.33, *p* = 0.006; Morphology: adj R^2^ = 0.07, *p* = 0.14). Finally, a marginally significant relationship (adj. R^2^ adj. = 0.12, *p* = 0.08) was observed between richness derived from the whole metagenome dataset and specific conductivity.

**Figure 6.**
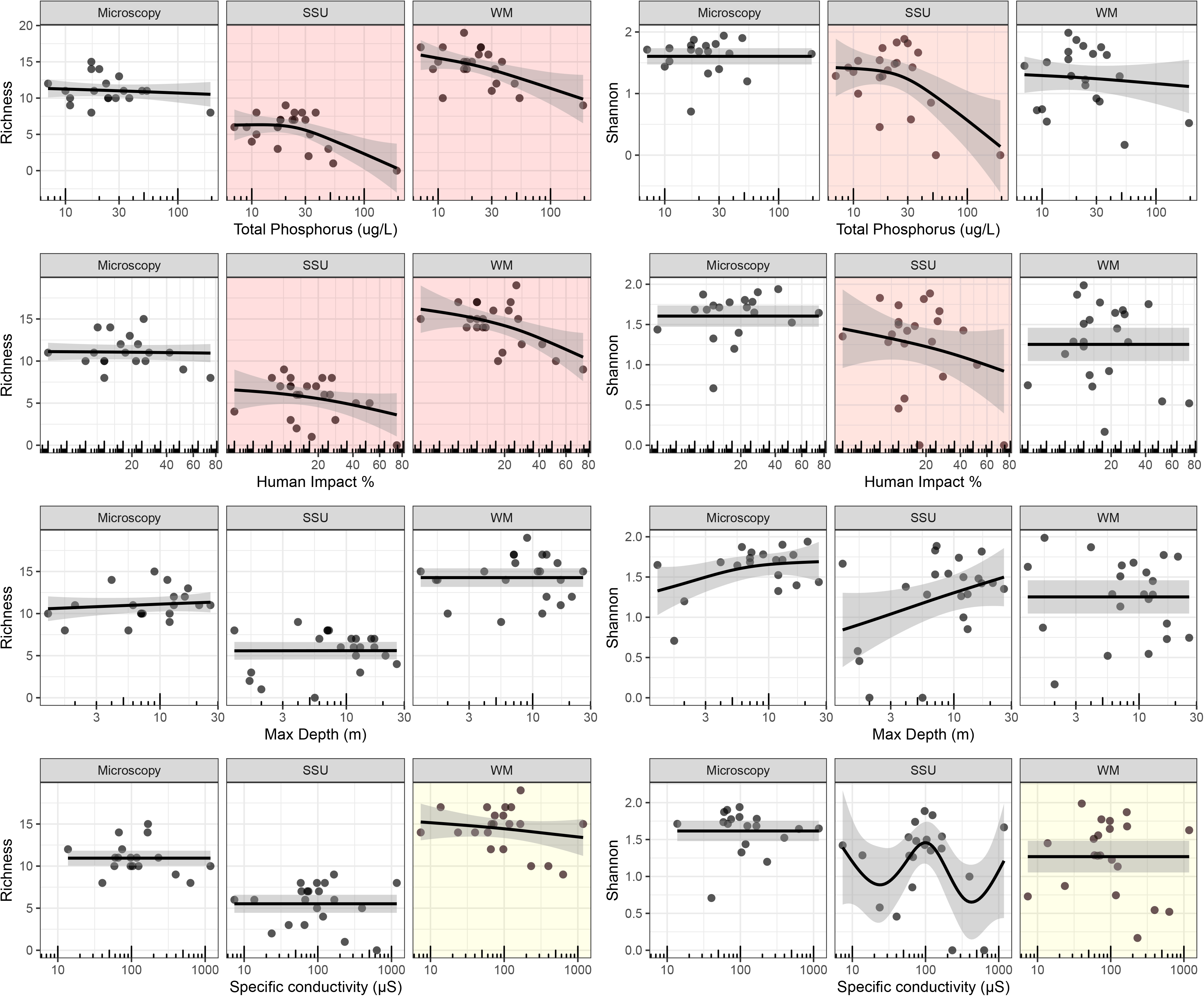
Diversity metrics (left: Taxonomic richness, right: Shannon entropy) estimated from microscopy, SSU rRNA genes (SSU), and whole metagenome (WM) datasets plotted against environmental gradients using generalized additive model (GAM). Environmental data were log transformed, except for the human impact index (expressed as percentage), which was arcsine transformed. Red and yellow backgrounds identify significant and marginally significant relationships, respectively. Adjusted R-squared and *p*-values for each GAM are listed in Supplementary Table S2)

For Shannon diversity, a significant fit was found between the SSU rRNA gene data and TP (adj. R^2^ = 0.18, *p* = 0.03) and between SSU rRNA data and human impact index (adj. R^2^ = 0.15, *p* = 0.056). The fit between Shannon diversity derived from whole metagenome data and specific conductivity was marginally significant (adj. R^2^ = 0.11 *p* = 0.098) and no significant relationships emerged with lake depth (**Figure 6 & Supplementary Table S2**).

## 4. Discussion

Consistent with other comparative analyses between eDNA metagenomics and morphological approaches (Stat et al., 2017; Singer et al., 2020), our results show that the match is not perfect. Overall, we detected modest congruence in taxon relative abundance across platforms and varying levels of congruence between analytical platforms when we considered presence-absence data. Interestingly, diversity metrics across all analytical platforms showed similar responses to epilimnetic phosphorus concentration, which is often considered a limiting nutrient in many lakes in Eastern Canada. Many important improvements can be implemented in future metagenomic work to help refine the robustness of this approach applied to metazoan biodiversity eDNA surveys (section 4.3).

### 4.1. To what extent do water metagenomes represent zooplankton biodiversity?

Using a variety of statistical approaches, we found that zooplankton communities surveyed using morphological counts and metagenomic analyses were, at best, moderately correlated. While local diversity metrics were similar across platforms, whole metagenome analysis detected the highest richness of zooplankton taxa. It is also informative to compare the strength of our results with other eDNA – morphological comparisons. For example, Keck et al. (2021) conducted a meta-analysis of comparative metabarcoding and morphological studies, and found that eDNA detected significantly more taxa than morphological counts, as eDNA may contain traces of taxa distributed outside of the immediate sampling area. Although no such synthetic analysis has been done from metagenomes, we would expect a similar finding. In our molecular dataset, we found four Families of zooplankton that were not recorded as part of the morphological survey, but these taxa are either generally characterized as benthic or littoral associated so may not have been present as individuals in the immediate sampling area. For instance, the Bdelloid rotifers *Adinetidae* and *Philodinidae,* which we only found via the whole metagenome analysis, are typically found to live on plants or debris in waters with dense vegetation and are generally not caught in plankton tows (Wallace and Snell, 2010). Across the metabarcoding and metagenomic literature, many have argued that eDNA approaches are more complementary to morphological approaches rather than directly exchangeable, and the coherence between metagenomic and metabarcoding for eukaryotic diversity surveys needs further detailed investigation (Garlapati et al., 2019; Cordier et al., 2020).

### 4.2. Shotgun sequencing reveals diversity patterns over broad environmental gradients

To explore diversity patterns over broad environmental gradients and among analytical platforms, richness and Shannon diversity metrics were plotted against epilimnetic total phosphorus (TP), specific conductivity, lake depth, and human impact index estimated in the 22 lakes (Huot et al., 2019). Based on our preliminary dataset, we found relatively consistent patterns in zooplankton diversity across analytical platforms, indicating that shotgun sequencing shows promise for investigating ecological gradients in freshwater systems. Our findings are consistent with results reported by Singer et al. (2020) from a marine system, where despite revealing contrasting taxonomic diversity, both the metagenomic and metabarcoding data revealed similar ecological patterns, which in turn were useful to infer factors related to the ecosystem health and function.

### 4.3. Limitations to the metagenomic approach and prospects

Limitations of eDNA-based approaches have been widely studied, although these have been primarily based on PCR-based approaches. Challenges relate mostly to the availability and quality of eDNA itself in water, whereby investigators have identified the conditions contributing to eDNA degradation (Barnes et al., 2014) or the transport of eDNA over long distances (Deiner and Altermatt, 2014). Although our knowledge of these factors is constantly improving the robustness of eDNA research, there are also other aspects of the workflow - from sampling strategies to bioinformatics – which need to be improved to strengthen metagenomic approaches applied to the study of metazoans in the environment.

#### 4.3.1. Methodological considerations

Our metagenomes were produced for the primary purpose of examining bacteria and archaea and thus the volume of water filtered was only ~250 ml to 500 ml, depending on how much water could be passed through the 0.22 μm membrane before it clogged (Garner et al., 2020). In contrast, the morphological identification of zooplankton was performed on tow haul samples collected from tens of liters across the full water column that were then concentrated to a few hundred milliliters. Furthermore, because the samples for DNA analysis were collected over the photic zone only, they might not fully represent the rotifer and crustacean zooplankton samples which were collected from below the thermocline and over the full water column, respectively. Overall, we are looking at diverging sampling efforts and distributions across approaches, which may have brought about some of the differences.

Filter size selection could also play an influential role and may allow one to filter more water in future studies. We opted for a size selection step where we first pre-filtered water through a 100 μm mesh, which selected mostly for extra-organismal DNA (i.e. DNA that is no longer found within an organism, as opposed to organismal DNA) (Rodriguez-Ezpeleta et al., 2021). It is however likely that gametes and other juvenile stages in cladocerans and copepods passed through the 100 μm mesh and got caught on the 0.22 μm membrane, which may have contributed to the inflated number of reads assigned mainly to *Daphniidae* and copepods in the whole metagenome dataset. Selecting a filter with a pore size better suited to our target organisms may lead to a better overall coverage by minimizing the allocation of sequencing effort to DNA from non-target microbial taxa. For instance, 0.45 μm cellulose nitrate filters have been shown to yield consistent results for fish metabarcoding with high repeatability between filtration replicates (Li et al., 2018). Type of filter, pre-filtration step and pore size have all been identified as factors determining the final yield of eDNA, with differences observed between taxa and systems (Bowers et al., 2021)

Filtering larger volumes of water combined with an increased sequencing depth may help yield a higher number of reads and more diversity for Eukarya, which are very much underrepresented in metagenomes in contrast with bacteria and archaea. Similar to earlier research, we found that the proportion of recovered eukaryotes tends to be < 0.5% of the total read assignments, either with a genome wide approach (Stat et al., 2017; Cowart et al., 2018) or a gene-centric approach (Tedersoo et al., 2015). In contrast, the filtration of 10 L of water targeting extracellular DNA combined with ultra-deep sequencing was shown to yield ~100 million reads per metagenome from a brackish lagoon and improved the coverage for Eukarya to a proportion corresponding to over 4% of the total number of reads (Manu, 2021). Other emerging target enrichment techniques such as hybridization capture have great potential to improve the detection of metazoans in metagenomes (Seeber et al., 2019; Sevigny et al., 2021). Hybridization capture utilizes RNA probes carefully designed to bind the gene region of interest, enhancing the signal of desired taxa without introducing PCR-induced biases. Recent results based on ultra-deep sequencing have shown that the coverage for eukaryotes may be improved when combining shotgun sequencing with DNA target-capture methods (Manu, 2021). Alternatively, metatranscriptomics is an emerging and promising approach for characterizing zooplankton communities. A recent study by Lopez et al. (2021) comparing zooplankton estimates from observational with both amplicon sequencing and metatranscriptomics datasets has revealed higher congruence of observational zooplankton abundance and composition with metatranscriptomics estimates compared to amplicons sequencing using genomic (gDNA) and complementary DNA (cDNA) amplicons sequencing.

#### 4.3.2. Bioinformatics considerations

Carefully designed bioinformatic workflows are crucial for robust taxonomic assignment of sequencing reads. Our data suggest that using the whole metagenomic reads can capture the widest pool of biodiversity, but that the taxonomically informative gene markers, such as the SSU rRNA genes in eukaryotes better reflected the observed relative abundance of zooplankton families based on microscopy. We also found that in using a targeted approach using SSU rRNA genes as taxonomic markers, several taxa were missing or did not get taxonomically assigned using the lowest common ancestor (LCA) algorithm, even though they were present in relatively high abundances in microscopy counts. Some taxa were consistently missing or almost absent from our genetic datasets (such as the *Bosminidae*), despite being one of the most abundant taxonomic groups in microscopy. Such incongruences between traditional and metabarcoding data have frequently been reported (see Keck et al., 2021). We hypothesize that part of the issue with missing taxa in our metagenomes is caused by the same bioinformatic limitations as in any genetic-based study: the current lack of complete reference molecular data limits our ability to assign taxonomy to sequence reads. In molecular datasets, and especially in shotgun sequencing data, a large portion of the reads generated only get assigned to Class level or lower. These reads are typically filtered out bioinformatically, meaning that they are not considered in the estimation of diversity indices or in comparisons with other datasets. Therefore, we might be widely underestimating the abundance and diversity of certain taxonomic groups which are not populated in reference databases.

Since using the full metagenome read set is computationally intensive and does not appear to yield a higher correlation with morphology-based identifications, a reasonable compromise that might increase coverage without multiplying computation efforts could be the combination of a few targeted genetic markers. For example, metagenome reads mapped against both the SSU and LSU rRNA gene markers has been shown to improve taxa recovery in a study of marine plankton from DNA preserved in marine sediment (Armbrecht, 2020). In addition to nuclear SSU rRNA genes, we investigated the mitochondrial cytochrome c oxidase subunit I (COI) but found the coverage for this marker to be very low for metazoans, most likely due to the generally lower cellular abundance of mitochondria compared to ribosomes. For this reason, we did not pursue further metagenomic COI marker analyses.

There is clearly an urgent need for curated molecular databases to improve interpretation of eDNA-based molecular datasets. This is especially the case for freshwaters, where monitoring efforts are limited and yet provide habitat for a disproportionate number of taxa per unit area (Strayer and Dudgeon, 2010). Currently there are insufficient data for many taxa such that we cannot even assess the state of ~40% of freshwater species in Canada (Desforges et al., 2021). Initiatives to improve sequencing coverage of eukaryotic biodiversity are underway, including the Barcode of Life Data System (BOLD) (Ratnasingham and Hebert 2007), the International Barcode of Life (IBOL) (https://ibol.org/), the Earth BioGenome Project (https://www.earthbiogenome.org/), i5K for arthropods (Robinson et al., 2011), and Diat.barcode for Diatoms (Rimet et al 2019). Such initiatives will multiply the number of curated references for taxonomic marker genes, which is key to improving taxonomic assignments in eDNA studies.

## 5. Conclusion

In this study, metagenomics and classical morphological analyses of zooplankton applied to 22 freshwater lakes yielded contrasting abundance estimates but comparable diversity assessments at the family level. Metagenomics detected more taxa, including some that generally live outside the pelagic photic zone where the samples were taken, which is to be expected given the persistence and transport of eDNA in nature. Although metagenomic techniques still need to be improved with better adapted sampling protocols and refined bioinformatics pipelines specific to eukaryotic genomes, our results suggest enormous potential for extending metagenome analysis to the investigation of zooplankton and other aquatic micro- and macro-eukaryotes. Our comparative study contributes to a better understanding of how the metagenomic approach might contribute to biodiversity and ecological assessments in complement to other traditional and eDNA approaches.

## Acknowledgements

We thank the NSERC-funded LakePulse Network (2017-2019), Yannick Huot for leading LakePulse, Marie-Pierre Varin and Maxime Fradette for coordinating sampling and lab work. We are grateful to Vera Onana (Concordia University) for her contribution to producing some of the molecular data. We additionally thank Cindy Paquette (UQAM) for providing curated zooplankton identification and count tables, and Michelle Gros (McGill University) for the rotifer identification and help compiling zooplankton data. We also wish to acknowledge the contribution of the high-throughput sequencing platform of the McGill University and Genome Québec Innovation Centre (Montreal, Canada). M-ÈM received a postdoctoral fellowship co-funded by the Groupe de recherche interuniversitaire en Limnologie (GRIL), Liber Ero, and McGill University. The project was supported by LakePulse. IG-E, DAW and MEC acknowledge additional funding from the Canada Research Chairs program and NSERC Discovery program. The authors declare no conflict of interest.

## Data archiving statement

Sequence data were submitted to the EBI metagenomics platform for analysis and archiving under Study MGYS00003941.

