## Supplementary Materials for "Comparative analysis of zooplankton diversity in freshwaters: What can we gain from metagenomic analysis?"

**Supplementary Table S1.** Sites name, locational coordinates, and summary of relevant lake and sample characteristics.

| **Lake name** | **LakePulse ID** | **Latitude** | **Longitude** | **Ecozone** | **Size class** | **Human Impact index (%)** | **[TP] epilimnion (µg/ml)** | **Trophic status*** | **Specific conductivity (µS)** | **Lake max. depth (m)** | **Type of data** |
| --- | --- | --- | --- | --- | --- | --- | --- | --- | --- | --- | --- |
| Lac Parenthèses | LP06-071 | 50.379073 | -68.804204 | Boreal Shield | medium | 12 | 9 | Oligotrophic | 0.006 | 17.0 | WSM |
| Lac Simoncouche | LP06-095 | 48.232562 | -71.249643 | Boreal Shield | medium | 5 | 24 | Mesotrophic | 0.110 | 7.0 | WSM |
| Lac Forget | LP06-102 | 50.718456 | -71.765768 | Boreal Shield | medium | 24 | 7 | Oligotrophic | 0.011 | 13.0 | WSM |
| Lac Paula | LP06-126 | 48.99124 | -74.02811 | Boreal Shield | medium | 19 | 30 | Mesotrophic | NA | 17.0 | WSM |
| Lac Duminy | LP06-127 | 48.987717 | -74.06735 | Boreal Shield | small | 11 | 32 | Eutrophic | 0.019 | 1.6 | WSM(noR) |
| Lac Ouellette | LP06-136 | 46.720459 | -75.437217 | Boreal Shield | medium | 31 | 48 | Eutrophic | 0.060 | 13.0 | WSM |
| Wabun Lake | LP06-156 | 45.225314 | -76.83351 | Boreal Shield | small | 0 | 10 | Oligotrophic | 0.105 | 26.0 | WSM |
| Fraser Lake | LP06-161 | 45.191424 | -77.646029 | Boreal Shield | medium | 15 | 23 | Mesotrophic | 0.065 | 16.0 | SM |
| Bright Lake | LP06-220 | 45.191424 | -77.646029 | Boreal Shield | medium | 15 | 23 | Mesotrophic | 0.068 | 11.0 | WSM |
| Ritchie Lake | LP07-029 | 45.415706 | -65.96754 | Boreal Shield | small | 52 | 11 | Mesotrophic | 0.384 | 12.0 | WSM |
| Napadogan Lake | LP07-033 | 46.415881 | -66.943059 | Atlantic Maritime | small | 9 | 17 | Mesotrophic | 0.037 | 1.7 | SM |
| Lac Saint-Augustin | LP08-097 | 46.751029 | -71.390073 | Mixedwood Plains | medium | 77 | 195 | Eutrophic | 0.575 | 5.5 | WSM |
| Lac des Chicots | LP08-120 | 46.796823 | -72.522866 | Mixedwood Plains | medium | 30 | 33 | Eutrophic | 0.091 | 21.0 | WSM |
| Loch Garry | LP08-134 | 45.257686 | -74.702097 | Mixedwood Plains | medium | 7 | 20 | Mesotrophic | 0.141 | 4.0 | WSM |
| Lac Echo | LP08-138 | 45.892125 | -74.028641 | Mixedwood Plains | medium | 23 | 17 | Mesotrophic | 0.145 | 9.0 | WSM |
| Lac McCord | LP08-219 | 45.693927 | -76.46573 | Mixedwood Plains | small | 9 | 24 | Mesotrophic | 0.088 | 12.0 | WSM |
| Lac Volet | LP17-090 | 46.136558 | -70.809503 | Atlantic Highlands | small | 2 | 18 | Mesotrophic | 0.056 | 6.0 | WSM |
| Lac Fortin | LP17-091 | 46.119508 | -70.853652 | Atlantic Highlands | medium | 11 | 17 | Mesotrophic | 0.061 | 11.5 | WSM |
| Lac à la Truite | LP17-098 | 46.084681 | -71.507629 | Atlantic Highlands | medium | 17 | 53 | Eutrophic | 0.204 | 2.0 | WSM |
| Etang Burbank | LP17-107 | 45.778379 | -72.003987 | Atlantic Highlands | small | 28 | 37 | Eutrophic | 1.065 | 1.2 | WSM |
| Lac des Français | LP17-112 | 45.439487 | -72.22285 | Atlantic Highlands | small | 15 | 11 | Mesotrophic | 0.053 | 7.0 | WSM |
| Spooner Pond | LP17-113 | 45.738907 | -72.145011 | Atlantic Highlands | small | 22 | 28 | Mesotrophic | 0.089 | 7.2 | WSM |

*Trophic status is based on phosphorus concentration: oligotrophic (less than 10 µg/L), mesotrophic (10 – 30 µg/L) and eutrophic (greater than 30 µg/L). Type of data: W; whole mategenome, S; small subunit (SSU) rRNA genes, D; zooplankton microscopy, noR; no rotifer data.

**Supplementary Table S2.** R-squared and p-values for each generalized additive model (GAM; Figure 6) showing the relationship between diversity indices (richness, Shannon diversity) and environmental gradients. (Significant relationships are shown in bold and marginally significant relationships are shown in italics)

| Response variable | Type of data (analytical platform) | Explanatory variable | Data transformation | R-squared adjusted | p-value |
| --- | --- | --- | --- | --- | --- |
| Taxonomic richness | Microscopy | Total phosphorus(µg/L) | Log+1 | 0.03 | 0.24 |
|  | SSU |  |  | **0.21** | **0.03** |
|  | WM |  |  | **0.29** | **0.01** |
| Shannon entropy | Microscopy |  |  | <0.1 | 0.86 |
|  | SSU |  |  | **0.18** | **0.03** |
|  | WM |  |  | 0.05 | 0.18 |
| Taxonomic richness | Microscopy | Maximum depth (m) | None | 0.002 | 0.32 |
|  | SSU |  |  | <0.1 | 0.96 |
|  | WM |  |  | <0.1 | 0.98 |
| Shannon entropy | Microscopy |  |  | 0.002 | 0.32 |
|  | SSU |  |  | 0.05 | 0.19 |
|  | WM |  |  | <0.1 | 0.71 |
| Taxonomic richness | Microscopy | Specific conductivity (µS) | Log+1 | <0.1 | 0.38 |
|  | SSU |  |  | <0.1 | 0.54 |
|  | WM |  |  | *0.12* | *0.08* |
| Shannon entropy | Microscopy |  |  | <0.1 | 0.50 |
|  | SSU |  |  | <0.1 | 0.50 |
|  | WM |  |  | 0.11 | 0.98 |
| Taxonomic richness | Microscopy | Human impact index (%) | Arcsine | 0.07 | 0.14 |
|  | SSU |  |  | **0.23** | **0.022** |
|  | WM |  |  | **0.33** | **0.006** |
| Shannon entropy | Microscopy |  |  | <0.1 | 0.42 |
|  | SSU |  |  | **0.15** | **0.056** |
|  | WM |  |  | 0.06 | 0.156 |

**Supplementary File 1.**

To verify if the crustacean zooplankton and rotifer counts were consistent (e.g., without major discrepancies due to the slight difference in the integrated water sampling depth, or due to degradation of samples over time, as the rotifers were counted a few years after the crustacean zooplankton samples), we picked an easily identifiable zooplankton group (the *Bosminidae*, comprised of *Eubosmina* or *Bosmina* spp.) to count in parallel with the rotifers for comparison with the original zooplankton dataset. The new *Bosminidae* counts confirmed that the preserved samples for rotifer counting were representative of the original zooplankton samples (adj. R^2^ = 0.73 F-statistic: 51.5 on 1 and 18 DF, *p =* 1.111e-06; Supplementary Figure S1).

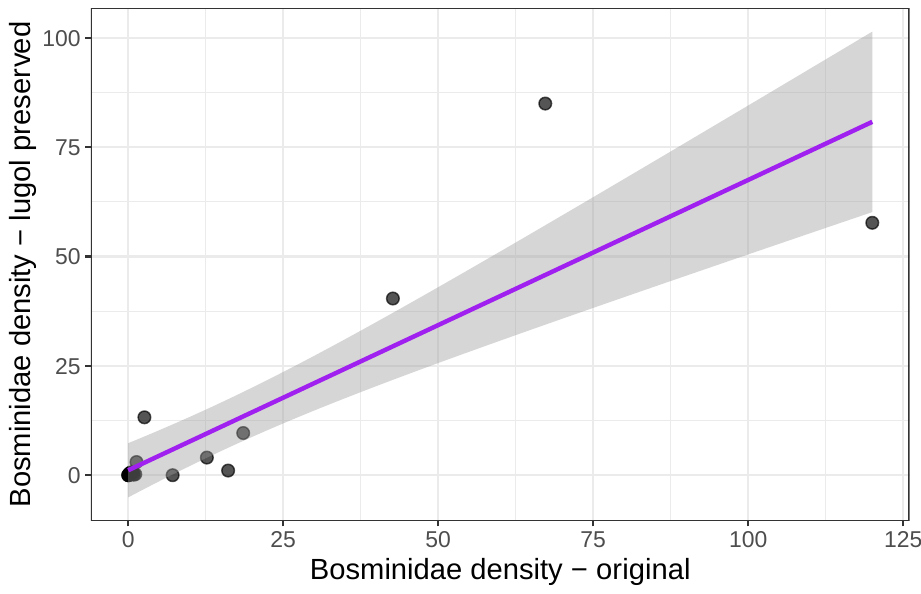

**Figure S1.** Least-square regression plot showing the positive relationship between original *Bosminidae* species counts and lugol-preserved *Bosminidae* counts confirming that the samples used for rotifer identification were well-preserved and were reflecting the original cladoceran zooplankton densities.

**Supplementary File 2**

Detailed bioinformatics workflow for the metagenomic approaches based on whole metagenome and SSU rRNA gene subset.

**Step 1**: Preprocessing Illumina NovaSeq (6000 S4 PE 150) raw reads using Trimmomatic v0.38 (Bolger et al., 2014)

java -jar trimmomatic-0.38.jar PE R1.fastq.gz R2.fastq.gz forward_paired_R1.fq.gz forward_unpaired_R1.fq.gz reverse_paired_R2.fq.gz reverse_unpaired_R2.fq.gz ILLUMINACLIP:NovaSeq_seq_adaptor_PE.fa:2:30:10 LEADING:3 TRAILING:3 SLIDINGWINDOW:4:15 MINLEN:36

**Step 2**: Quality check using FastQC (Andrews 2010)

**Step 3**: a) Whole metagenome approach: Merge forward and reverse reads using PEAR 0.9.10 (Zhang et al., 2014)

pear -f R1.fq -r R2.fq -y 32G -j 12 -p 0.01

b) SSU subset gene prediction approach: Workflow of EBI MGnify (available on EBI website: https://emg-docs.readthedocs.io/en/latest/analysis.html#raw-reads-analysis-pipeline)

**Step 4**: Read mapping to nucleotide database populated with all eukaryote sequences in the NCBI nt database:

4.1. convert fasta database to BLAST database:

makeblastdb -parse_seqids -in nt.$Eukaryote.fa -dbtype nucl -out nt.Eukaryote

4.2. Map reads against nucleotide database

blastn db nt.Eukaryote -evalue 0.001, min % ID 70, max number of hits 30

**Step 5**:	Lowest Common Ancestor (LCA) analysis using MEGAN (MEtaGenome ANanlyzer) (Huson et al. 2016):

Database file downloaded: megan-nucl-Jul2020.db.zip

LCA parameters: Min score 80; max expect 0.001, top percent 10.0, min support 2, min complexity filter 0.1

**Supplementary Figures**

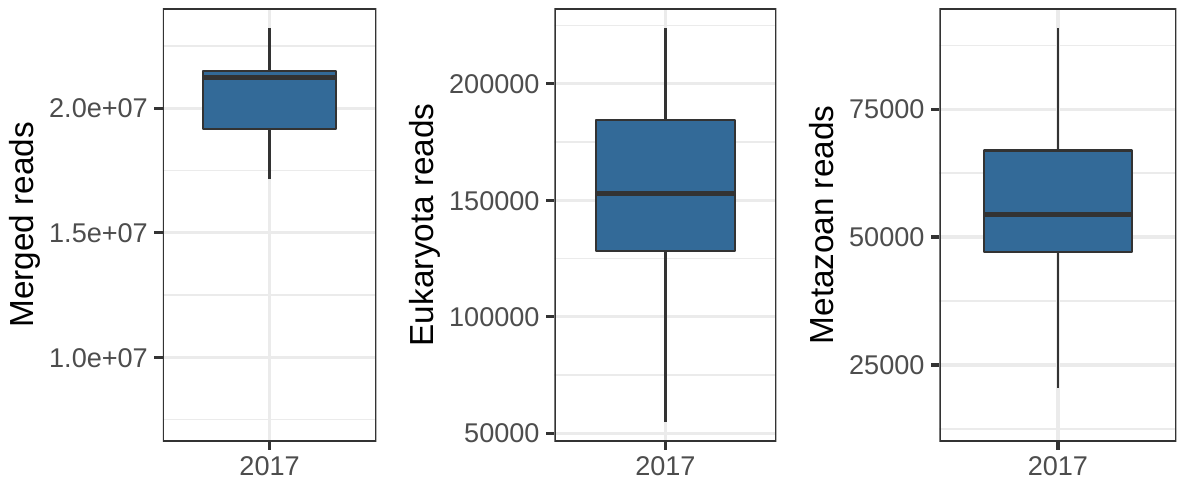

**Supplementary Figure S2.** Basic shotgun metagenome stats (number of quality-checked merged reads, reads assigned to eukaryote and reads assigned to metazoans). Merged reads assigned to metazoans generally accounted on average for 0.27% (max=0.47%) of the total merged reads count, and for 34% (max=46%) of reads assigned to Eukaryota.

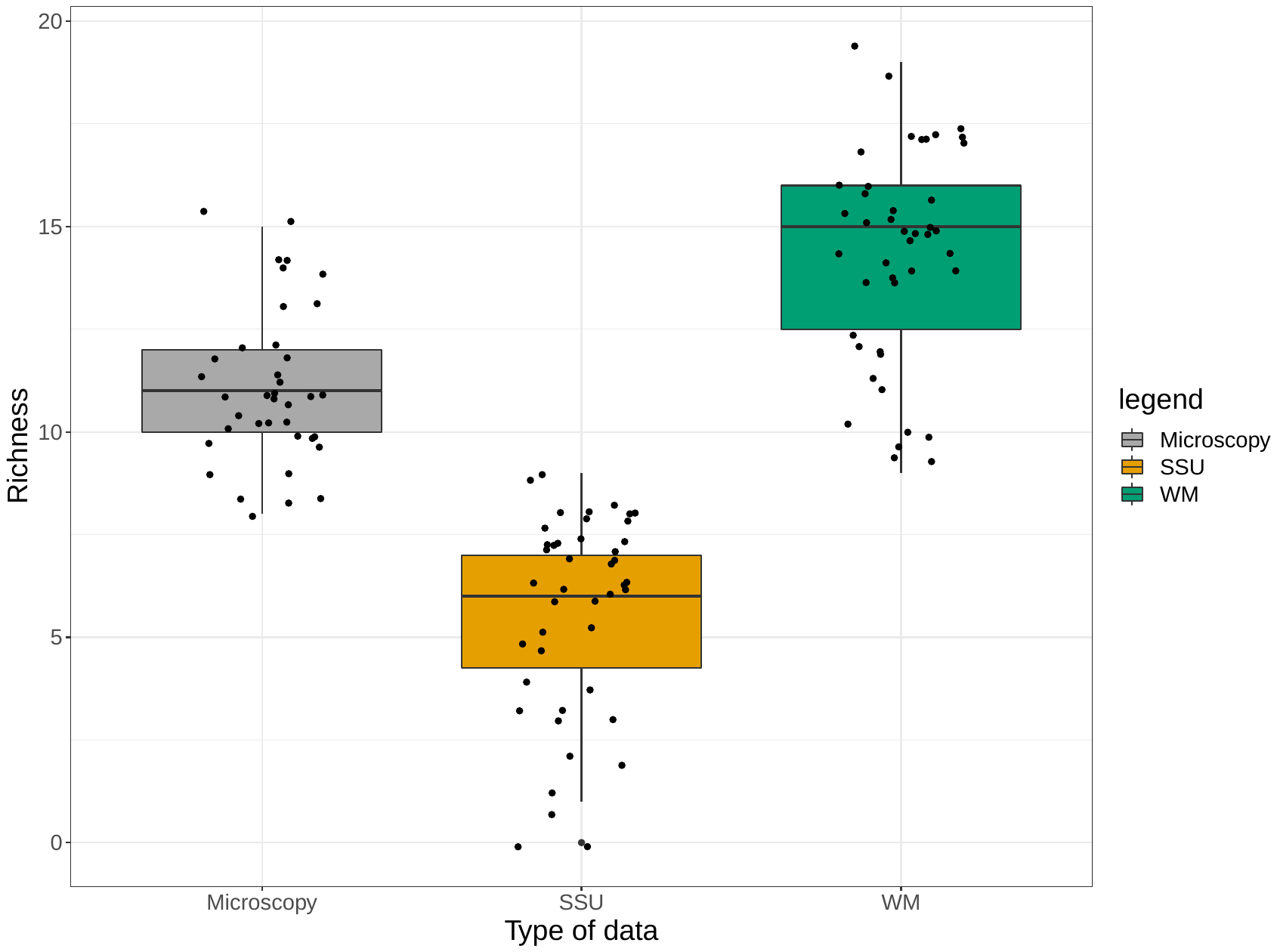

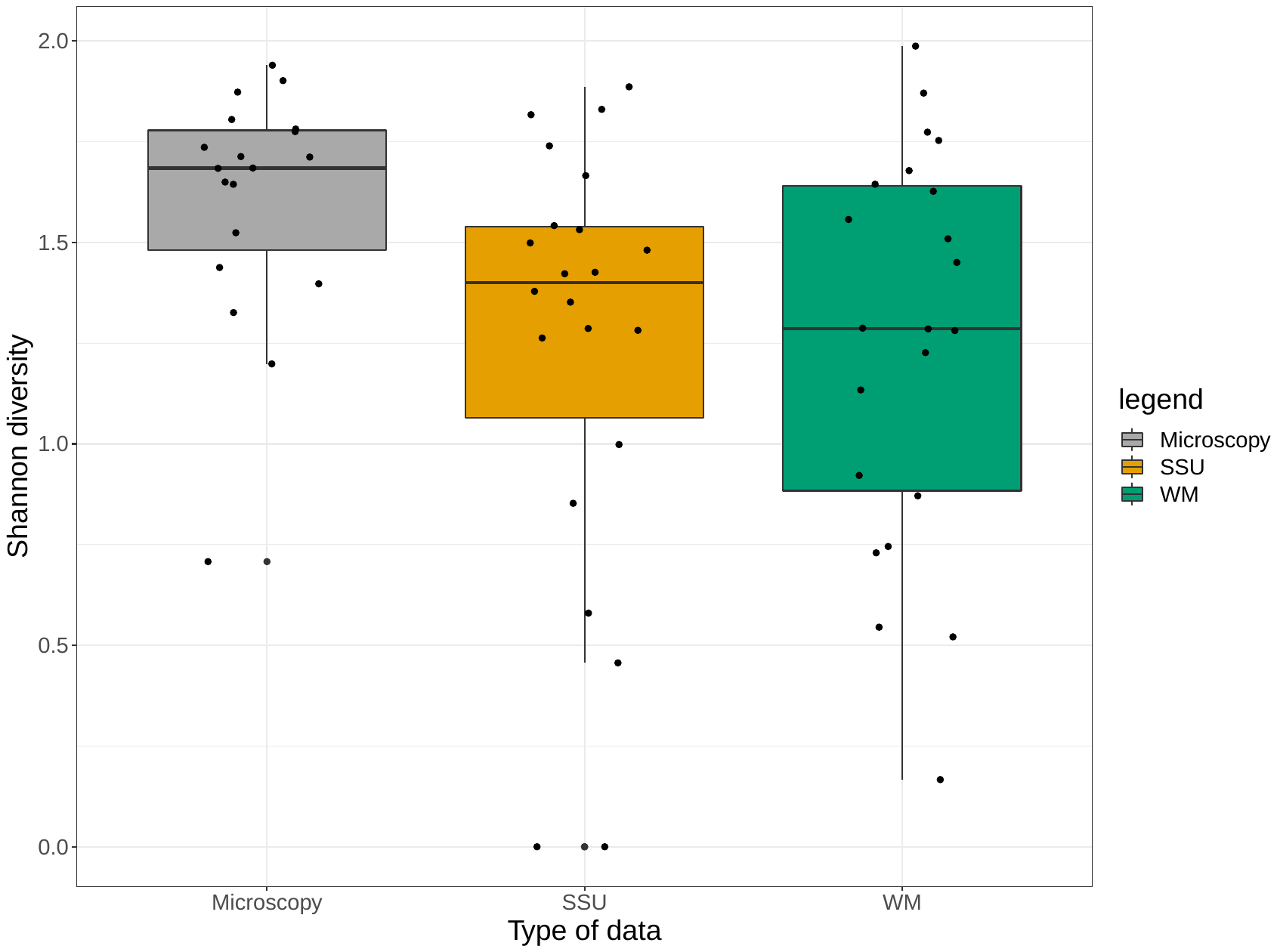

**Supplementary Figure S3.** Summary of the family-level taxonomic richness and Shannon diversity across the 22 lakes for each analytical platform (grey: microscopy, orange: small subunit subset, green: whole metagenome dataset.

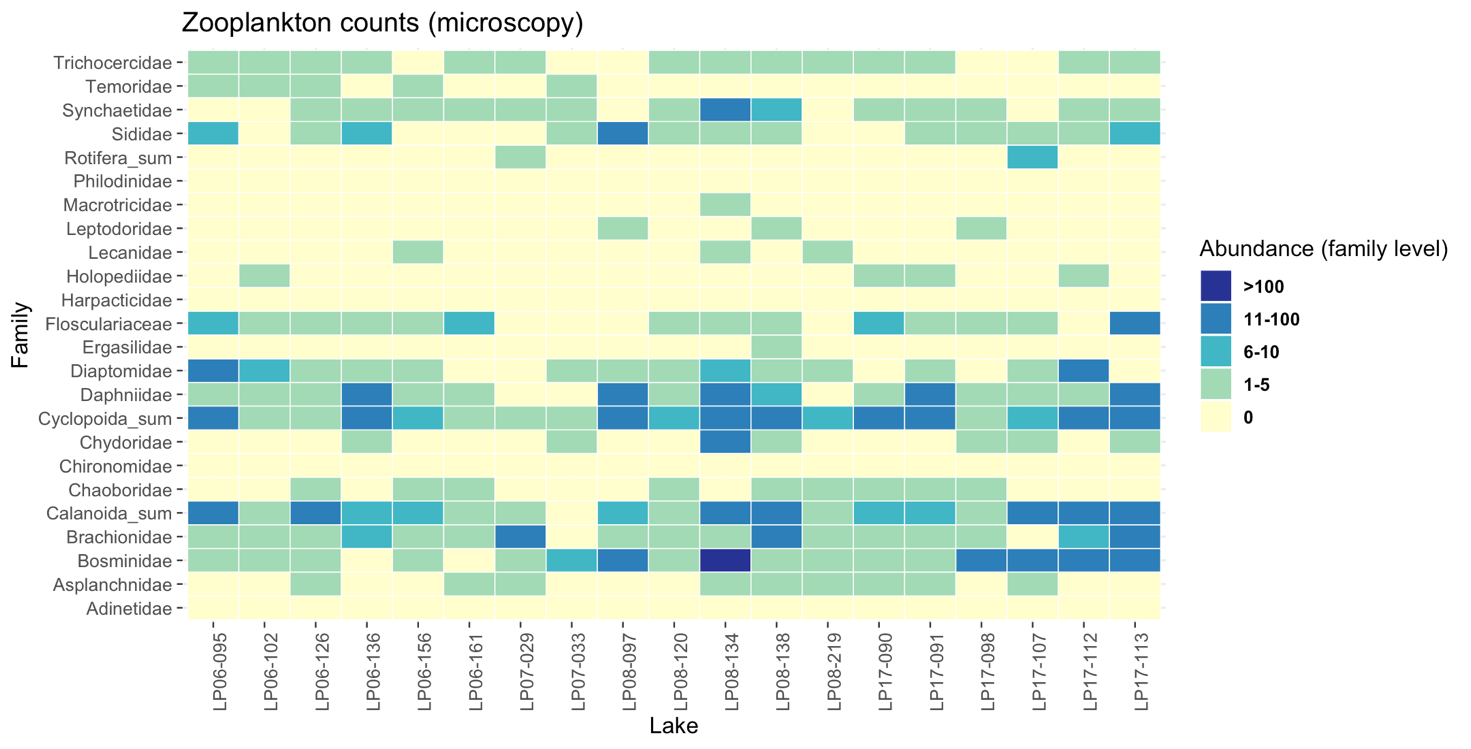

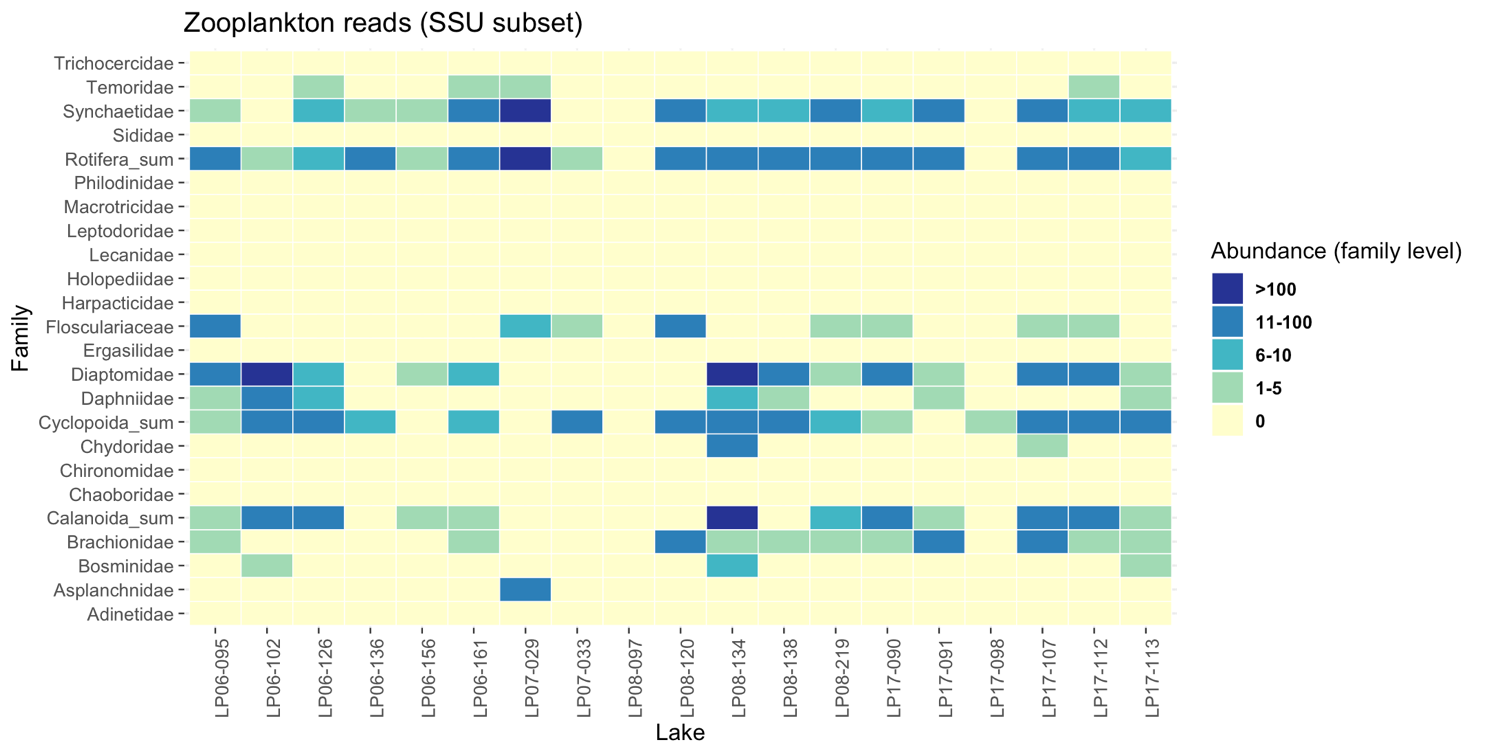

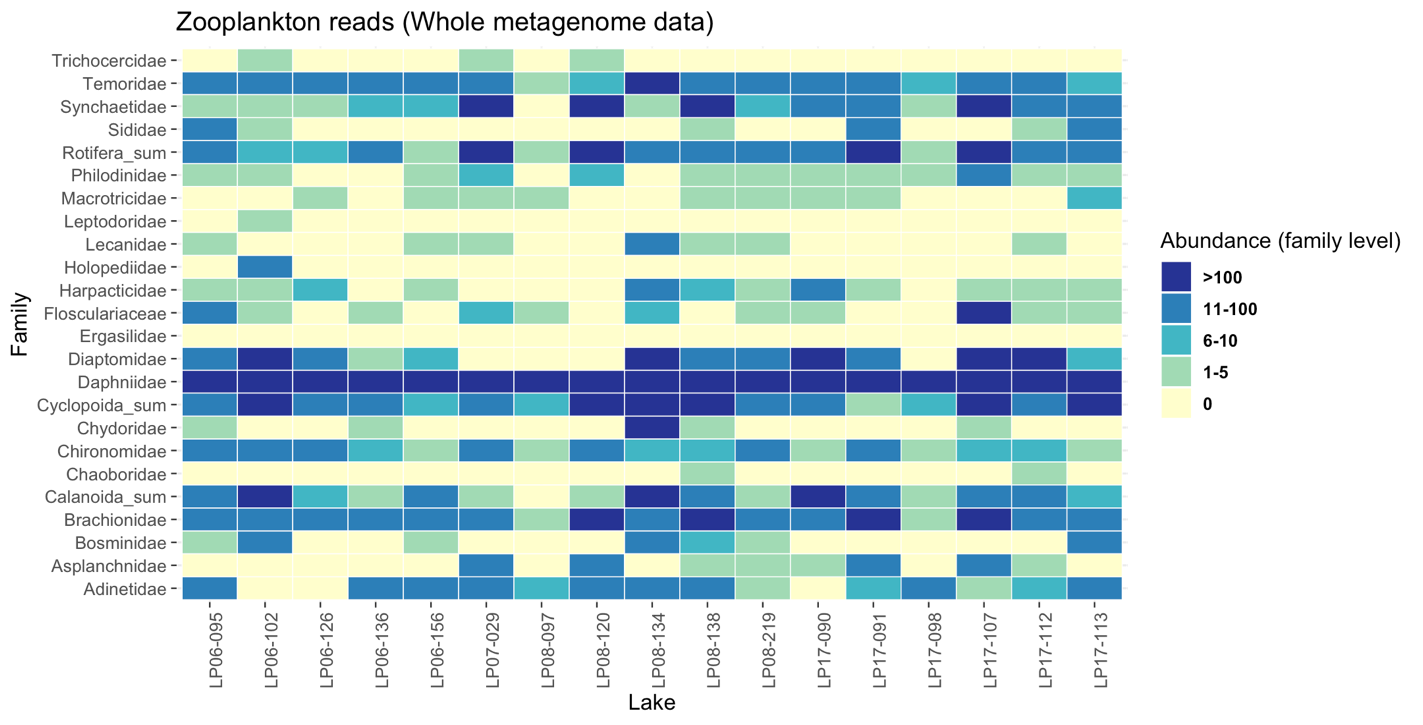

**Supplementary Figure S4.** Heatmaps showing the abundance of zooplankton families derived from each dataset of taxonomy density, whole metagenome, and small subunit (SSU) subset analysis.
